# Reference mapping and whole-genome alignment yield comparable phylogenomic inferences

**DOI:** 10.64898/2026.10.20.760391

**Authors:** Michael V. Westbury, David A. Duchene

## Abstract

Reference-genome choice can introduce mapping bias when sequencing data are analysed against a heterospecific genome, but it remains unclear how strongly this bias affects species-level phylogenomic inference. We tested this across Ursidae using nine species spanning approximately 20 million years of divergence, including the extinct giant short-faced bear (*Arctodus simus*). We independently mapped the same sequencing datasets to four reference genomes spanning a broad range of evolutionary distances and compared these with two assembly-based whole-genome-alignment approaches. We developed PhyloSlide to analyse all treatments consistently, offering a streamlined pipeline for sliding-window phylogenomics. Using this approach, we found that mapping reference choice strongly affected sequence recovery, with missing data generally increasing with evolutionary distance. Nonetheless, all six treatments recovered an identical species-tree topology and highly similar gene and site concordance factors. Relative divergence-time estimates were also largely consistent between approaches. Together, these findings demonstrate that mapping sequencing reads from multiple species to a shared reference genome can yield phylogenomic inferences comparable to whole-genome alignment, despite substantial reference-dependent sequence loss across millions of years of evolutionary divergence.

## Introduction

Reconstructing evolutionary relationships increasingly relies on genome-scale data, which provide thousands of loci distributed across the genome and allow phylogenetic signal and discordance to be examined at much finer resolution than individual genes or markers (Feng et al. 2020). However, generating comparable genomic datasets across multiple species remains challenging because high-quality reference genomes are not available for every taxon. This is particularly relevant when studying extinct species, which often lack a conspecific reference genome and for which generating a new high-quality assembly may be limited by the quantity and quality of surviving DNA (Liu et al. 2021). A common solution is therefore to map sequencing reads from multiple species to the genome of a related taxon and extract homologous regions from the resulting consensus sequences (Barnett et al. 2020; Sinding et al. 2026). This approach enables genome-scale phylogenetic analyses without requiring a *de novo* assembly for every species, but it may also introduce systematic bias if reads that differ from the reference are less likely to map successfully. As evolutionary distance between the sample and reference increases, this bias could progressively impact downstream analyses, including estimates of phylogenetic relationships and divergence times.

The consequences of reference bias are well established for several population-genomic analyses. Increasing phylogenetic distance between sample and reference can reduce sequence recovery and bias estimates of genetic diversity, runs of homozygosity, and demographic history (Prasad et al. 2022; Akopyan et al. 2025). Reference-genome choice and bioinformatic filtering can also influence phylogenetic topology, branch lengths, and support in SNP-based analyses (Rick et al. 2024; Mohn et al. 2026). Reference bias presents an additional challenge for ancient DNA, where short and damaged molecules are already more difficult to align accurately (Dolenz et al. 2024).

Assembly-based whole-genome alignment provides an alternative to reference mapping by identifying homologous regions directly among assembled genomes. However, this approach requires a suitable genome assembly for each taxon, which is not always available, particularly for extinct or poorly studied species, and computational requirements can increase substantially with genome size and taxon sampling (Armstrong et al. 2019). Consequently, mapping sequencing reads to an available reference remains an attractive and often necessary approach. Determining when reference mapping provides reliable phylogenomic inference is therefore important for comparative genomic studies.

Several tools have been developed to automate genome-scale phylogenetic inference, including sliding-window analyses (Hempel et al. 2022), assembly-based species-tree reconstruction (Gupta et al. 2025), and ortholog-based phylogenomics (Li et al. 2026). However, these approaches do not provide a unified workflow connecting sliding-window phylogenetic reconstruction with the assessment of genomic discordance and selection of suitable regions for divergence-time estimation. To address this, we developed PhyloSlide, an open-source pipeline that integrates genomic window extraction, phylogenetic inference, gene and site concordance analysis, and automated filtering of windows based on topological agreement, phylogenetic support, and clocklikeness. PhyloSlide also prepares alignments for downstream divergence-time estimation, supports transversion-only analyses of ancient DNA, and accommodates reference-aligned sequences generated through either read mapping or whole-genome alignment.

Here, we used Phyloslide to investigate the influence of reference-genome choice on phylogenomic inference across Ursidae, a mammalian family spanning approximately 20 million years of evolutionary divergence and including a rapid diversification of the six ursine bears within the last ~5 million years (Wooldridge et al. 2025). Using genomic data from all eight extant bear species and the extinct giant short-faced bear (*Arctodus simus*), we compared four reference-mapping treatments with two whole-genome-alignment approaches. We applied PhyloSlide consistently across all six treatments to determine whether reference-dependent differences in sequence recovery influence species-tree topology, genomic concordance, and relative divergence-time estimates.

## Results

Across 40 sample × reference combinations, mean sequencing depth ranged from 8.7× to 57.0×, and 80.9–98.6% of each reference was covered by at least one read (Table S1). Despite generally high coverage, reference choice strongly affected the amount of sequence retained in the final consensus genomes (Figure 1A).

**Fig 1.**
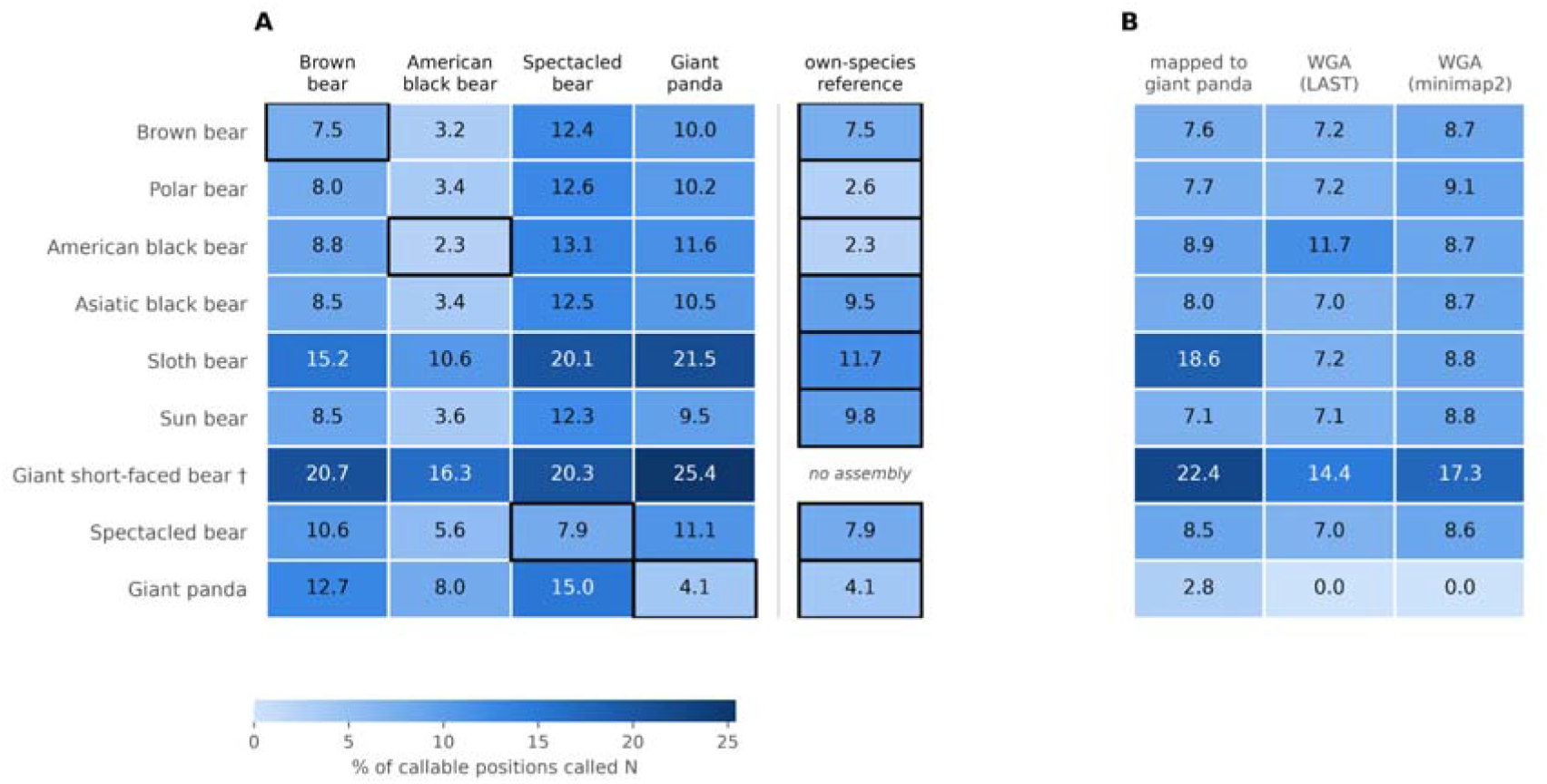
Missing data after read mapping and after whole-genome alignment. Each cell gives the percentage of callable reference positions emitted as N in the consensus sequence for that sample, where callable positions are those at which the reference assembly itself carries sequence. (**A**) Each of the nine samples mapped to each of four shared reference genomes (left) and to the reference genome of its own species (right). Cells outlined in black are those in which the reference genome is the sample’s own species. The giant short-faced bear has no assembly, so no own-species value exists. (**B**) Read mapping compared with both whole-genome alignment pipelines, all three measured over the same 24 giant panda anchor scaffolds (≥ 1 Mb, 94.2% of assembly length) on which the projected pseudo-genomes are defined. Giant panda values of 0.0 in the alignment columns arise because the anchor projected onto itself. Giant panda values differ between panels because A is scored over the complete assembly and B over the 24 scaffolds ≥1 Mb (94.2% of assembly length).

When samples were mapped to a conspecific assembly, 2.3–11.7% of callable positions were represented by N. Missing data generally increased with phylogenetic distance between the sample and reference. Across non-conspecific samples, median missingness was 4.6% using American black bear as the reference, 9.7% using brown bear, 10.8% using giant panda, and 12.8% using spectacled bear. The strongest effect occurred in the ancient giant short-faced bear sample, where missingness ranged from 16.3% against American black bear to 25.4% against giant panda.

We observed broadly similar sequence recovery when we compared read mapping with whole-genome alignment across the same 24 giant panda scaffolds (Figure 1B). Among well-covered modern samples, mapping reads to the giant-panda produced 7.1–8.9% missing sequence, compared with 7.0–7.2% for LAST and 8.6–9.1% for minimap2. LAST generally recovered slightly more sequence than minimap2. The larger differences observed for sloth bear and giant short-faced bear reflected dataset-specific properties, including shallow sequencing depth and the use of a mapped consensus as the giant short-faced bear input to whole-genome alignment.

The two whole-genome-alignment pipelines differed substantially in computational cost. LAST required each query genome to be divided into 28–81 chunks to limit memory requirements and consumed 946 CPU-hours across eight genomes, compared with 21 CPU-hours for minimap2, which aligned each genome in a single job. This represents an approximately 45-fold difference in computational time. In comparison, read mapping and consensus generation across the 40 sample–reference combinations required 7,482 CPU-hours.

### Topological inferences

Differences in sequence recovery had little effect on the number of windows available for phylogenomic analysis. Between 2,053 and 2,147 windows produced phylogenetic trees across the six treatments, and 270–307 windows passed the rigorous additional topology, support, and clocklikeness filters used for divergence-time estimation (Table S3). These windows produced alignments for molecular dating of 5.37–6.12 Mb in length.

All six treatments recovered the same ASTRAL species-tree topology (Figure 2A). Gene concordance factors (gCF) and likelihood-based site concordance factors (sCF) were also highly consistent among treatments, both varying by at most 3.6 percentage points for any given branch (Figure 2B,C). However, concordance varied substantially among branches and between measures. Crown Ursinae (node B) showed nearly complete gene concordance (gCF ~100%) but low site concordance (sCF ~35%), with relatively few decisive sites. In contrast, nodes C and F, involving *Ursus thibetanus, Melursus*, and *Helarctos*, showed low concordance for both genes and sites despite considerably more decisive sites, indicating substantial discordance among genomic regions. These patterns remained consistent across all six treatments.

**Fig 2.**
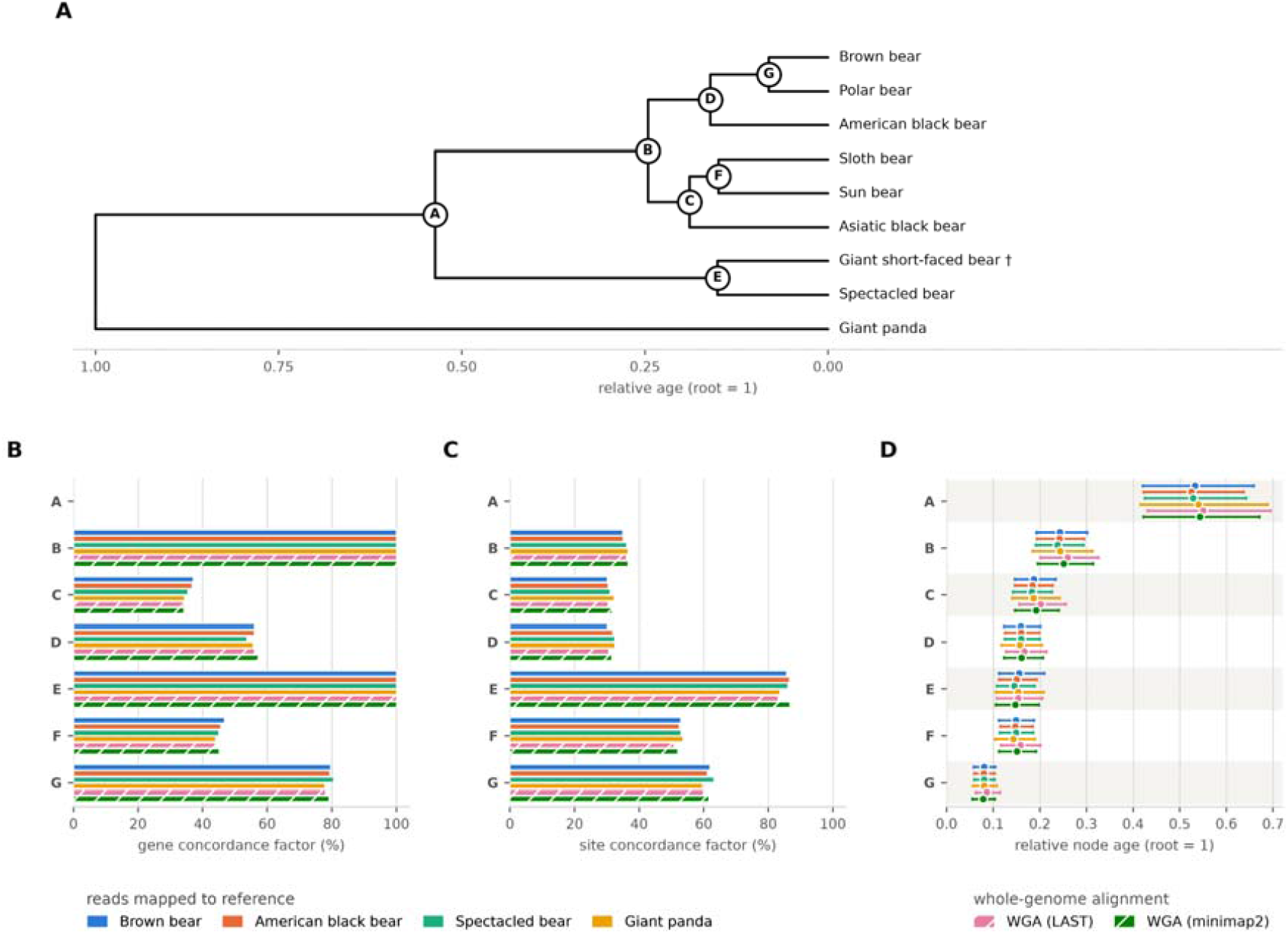
Species-tree topology, concordance factors, and relative divergence times. All four panels compare six analytical treatments: sequencing reads mapped to four reference genomes (brown bear, American black bear, spectacled bear, and giant panda) and two whole-genome-alignment approaches (LAST and minimap2), analysed using identical downstream phylogenomic parameters. (**A**) The shared ASTRAL species-tree topology, displayed as a relative timetree. The dagger indicates the extinct taxon. (**B, C**) Gene concordance factors (gCF) and likelihood-based site concordance factors (sCF), respectively, with one bar per treatment at each internal branch. Concordance factors are undefined for the outgroup-associated branch (node A), which is therefore left empty in both panels. (**D**) Relative divergence-time estimates obtained using MCMCtree under an independent-rates relaxed-clock model, with the root age fixed to 1. Points represent posterior means calculated from the pooled samples of two independent MCMC chains per treatment, and horizontal bars indicate the corresponding 95% highest posterior density intervals calculated directly from these pooled samples. HPD, highest posterior density; WGA, whole-genome alignment.

### Relative divergence times

Relative divergence-time estimates were similarly stable across treatments (Figure 2D). With the root fixed to 1, differences in the inferred timing of internal branches were negligible among treatments relative to the corresponding 95% highest posterior density intervals (HPDI). Specifically, the divergence time estimate of each internal node fell within the corresponding interval from all other treatments.

The two whole-genome-alignment treatments tended to produce slightly older ingroup divergence times than the reference-mapping treatments, with six of seven internal nodes shifting in the same direction. Comparing the two alignment approaches directly, minimap2 produced younger estimates than LAST at all seven nodes, with a median difference corresponding to 7.3% of the posterior credible-interval width.

## Discussion

Using PhyloSlide, a new pipeline developed to streamline sliding-window phylogenomic analyses, we found that substantial reference-dependent sequence loss had little effect on downstream phylogenomic inference. Although missing data increased with evolutionary distance between samples and reference genomes, all six analytical treatments recovered the same species-tree topology and highly similar gene and site concordance factors. Relative divergence-time estimates were also largely consistent across treatments.

Despite the substantial differences in sequence recovery, phylogenomic inference was remarkably stable across our treatments. Previous SNP-based studies have shown that reference-genome choice and bioinformatic filtering can alter inferred topology, branch lengths, support, and tree shape, particularly as evolutionary distance from the reference increases (Rick et al. 2024; Mohn et al. 2026). In contrast, our window-based analyses recovered an identical species-tree topology and highly qualitatively identical concordance factors across all six treatments. This suggests that the extent to which mapping bias propagates into phylogenetic inference depends strongly on the analytical framework. We reconstructed phylogenies from genome-wide consensus sequences distributed across thousands of genomic windows, excluded windows with excessive missingness, and summarised phylogenetic signals across loci rather than from a single SNP matrix. These features, implemented through our PhyloSlide pipeline, may reduce the influence of localised reference-dependent sequence loss.

The stability of concordance factors is particularly informative in Ursidae because gene-tree discordance is a known feature of bear evolution. Previous work has shown extensive incomplete lineage sorting and introgression among ursine bears, especially around relationships involving the Asian black, sun, and sloth bear lineages (Kutschera et al. 2014). More recent whole-genome analyses similarly recovered extensive gene-tree conflict despite strongly supported species trees and attributed much of this discordance to rapid diversification and ancient gene flow (Wooldridge et al. 2025). In our analyses, low site concordance did not always correspond to substantial conflict among genomic regions. At the crown Ursinae node (B), gene concordance was nearly complete (gCF ~100%), whereas site concordance was much lower (sCF ~35%), with relatively few decisive sites. In contrast, nodes C and F showed low gene and site concordance despite substantially more decisive sites, consistent with genuine genealogical discordance. These patterns remained consistent across all six treatments, supporting the interpretation that the observed genomic conflict reflects underlying evolutionary history rather than an artefact generated primarily by reference mapping.

Ancient DNA samples may be particularly susceptible to reference bias because DNA damage, short fragment lengths, and evolutionary divergence from available reference genomes can reduce mapping efficiency (Dolenz et al. 2024; Yuan et al. 2024). The giant short-faced bear combined these characteristics and showed the greatest reference-dependent sequence loss in our analyses. Nevertheless, its phylogenetic placement and concordance patterns remained consistent across all treatments. The only notable difference was a slightly younger estimated divergence from the spectacled bear when using the spectacled bear reference, consistent with potential reference attraction. However, this shift was small relative to posterior uncertainty, amounting to approximately 11% of the corresponding 95% highest posterior density interval width. Thus, even for the ancient sample most susceptible to reference-dependent sequence loss, the impact on downstream phylogenomic inference was limited.

In our analyses, whole-genome alignment offered little advantage over reference mapping, with both approaches recovering similar amounts of sequence, identical species-tree topologies, and highly consistent concordance patterns. However, the two whole-genome-alignment pipelines differed substantially in computational requirements, with minimap2 requiring approximately 45-fold fewer CPU-hours than LAST while recovering only slightly less sequence for most taxa. Despite their nearly identical topological and concordance results, the aligners produced small systematic differences in relative divergence-time estimates, indicating that alignment strategy can still influence downstream inference. Overall, our findings suggest that mapping sequencing reads to a shared reference genome provides a practical alternative to whole-genome alignment for within-family phylogenomics, particularly when suitable assemblies are unavailable.

Our study focused on a single mammalian family, predominantly high-coverage datasets, and one ancient taxon. Nevertheless, Ursidae provides a useful test case for within-family phylogenomics, encompassing substantial evolutionary divergence and relationships complicated by incomplete lineage sorting and introgression. Even at nodes with considerable genealogical discordance, reference-genome choice had little effect on inferred relationships or concordance factors. Our findings therefore suggest that mapping to a related reference genome can provide reliable phylogenomic inferences within a mammalian family, even when substantial reference-dependent sequence loss occurs. However, this robustness should not be assumed for comparisons across greater evolutionary distances or datasets with different characteristics, where reference-associated biases may have stronger effects. PhyloSlide provides a streamlined framework for assessing these effects when the suitability of a reference genome is uncertain.

## Methods

### Reference-genome benchmarking framework

To quantify the influence of reference-genome choice on phylogenomic inference, we designed a benchmarking framework in which we independently mapped the same set of ursid sequencing datasets to four reference genomes spanning different phylogenetic distances. We also generated two assembly-based whole-genome alignments to provide a comparison that did not rely on short-read mapping. We processed all six datasets using an identical phylogenomic workflow implemented in PhyloSlide, which we developed to standardise genomic window extraction, filtering, phylogenetic inference, species-tree estimation, concordance analysis, and selection of loci for divergence-time estimation.

### Samples and sequencing data

We assembled a nine-taxon panel comprising all eight extant ursid species and the extinct giant short-faced bear. We included the giant panda (*Ailuropoda melanoleuca*) as the outgroup. Table S1 provides sequencing accessions and library characteristics.

### Reference genome assemblies

We obtained eight ursid genome assemblies from NCBI (Table S2). We selected four as common benchmark references: brown bear (*Ursus arctos*), American black bear (*U. americanus*), spectacled bear (*Tremarctos ornatus*), and giant panda. We independently mapped every sequencing dataset to each of these four assemblies and, where available, additionally mapped each sample to a conspecific assembly.

We also used the giant panda assembly as the coordinate anchor for the whole-genome-alignment analyses. We restricted these analyses to its 24 scaffolds ≥1 Mb, which together contain 94.2% of the assembly sequence, and retained the soft masking distributed by NCBI.

### Read processing, mapping, and consensus generation

We processed and mapped reads using PlainMap v0.1 (Westbury 2026). We analysed modern libraries using the modern mode and the ancient giant short-faced bear dataset using ancient-fast. PlainMap used fastp v1.0.1 for read preprocessing (Chen et al. 2018), BWA v0.7.17-r1188 for alignment (Li and Durbin 2009), and SAMtools v1.13 for BAM processing (Li et al. 2009).

We generated consensus sequences using ANGSD v0.935 (Korneliussen et al. 2014) with - doFasta 2 -minQ 25 -minMapQ 25 -setMinDepthInd 5. ANGSD therefore emitted the most frequently observed nucleotide at sites supported by at least five reads passing base- and mapping-quality thresholds of 25 and emitted N at positions that failed these criteria.

We quantified missing sequence as the proportion of callable reference positions represented by N. We excluded reference assembly gaps but counted scaffolds receiving no sequence coverage as completely missing.

### Whole-genome alignment

We generated two independent assembly-based whole-genome-alignment datasets using LAST and minimap2. We used the 24 giant panda scaffolds ≥1 Mb as the anchor for both approaches.

For extant taxa, the genome assemblies generally represented different individuals from the resequencing datasets, while giant short-faced bear, for which no assembly is available, was represented by its spectacled-bear-mapped consensus sequence.

For the LAST analysis, we indexed the giant panda reference using lastdb -c to retain soft masking and aligned each query using LAST v1654 (Kiełbasa et al. 2011) with lastal -P1 -m 50 -E 0.05 -u 2, followed by last-split (Frith and Kawaguchi 2015). We converted the resulting MAF alignments to AXT format using maf-convert and generated chain files with axtChain -linearGap=medium.

We independently aligned the same query genomes using minimap2 v2.31 (Li 2021) with -cx asm10 --cs. We converted the resulting PAF alignments directly to UCSC chain format using paf2chain v0.1.1. Because paf2chain assigned PAF mapping-quality values as chain scores, these fell below the default minimum score of 2,000 required by chainNet. We therefore replaced each chain score with the total number of aligned bases in that chain.

Following chain generation, both approaches used the same UCSC chain/net workflow (v482): chainSort, chainNet, netSyntenic, netToAxt, and axtToMaf (Kent et al. 2002). We retained syntenic alignments and projected them onto giant panda coordinates to produce one pseudo-genome per taxon. We represented anchor positions without an aligned query base as N and discarded query insertions because they lacked an anchor coordinate.

### Sliding-window phylogenomic analyses with PhyloSlide

We analysed all four reference mapping datasets and both whole-genome-alignment datasets using PhyloSlide under identical downstream settings. PhyloSlide extracts homologous genomic windows from genome FASTA files, filters windows according to missing data and phylogenetic information content, infers window-specific phylogenies, and coordinates species-tree, concordance, and divergence-time analyses. PhyloSlide generates windows using BEDTools (Quinlan and Hall 2010) and extracts sequences using SAMtools faidx.

For this study, we sampled 20-kb windows at 1-Mb intervals across scaffolds ≥1 Mb. We retained windows when no taxon contained >50% missing sequence (--maxN 0.5) and the alignment contained at least 20 parsimony-informative sites (--minpi 20).

We inferred maximum-likelihood phylogenies independently for each retained window using IQ-TREE v2.4.0 (Bui Quang Minh et al. 2020) under GTR+R6 with 1,000 ultrafast bootstrap replicates (Hoang et al. 2018). The FreeRate model allows among-site rate categories to vary without constraining their distribution to a gamma model (Yang 1995; Soubrier et al. 2012).

We collapsed branches with <30% ultrafast bootstrap support before estimating species trees from the window trees using ASTRAL v5.7.8 (Sayyari and Mirarab 2016; Zhang et al. 2018). We rooted all species trees using the giant panda.

We calculated gene concordance factors (gCF) and likelihood-based site concordance factors (sCF) for each ASTRAL species-tree topology using IQ-TREE v2.4.0 (B. Q. Minh et al. 2020; Mo et al. 2023). We estimated sCF using 100 randomly sampled quartets per branch.

PhyloSlide also supports alternative window sizes and filtering thresholds, concatenated species-tree inference, and transversion-only analyses that mask A/G and C/T polymorphic columns when transition-associated damage is a concern. We did not apply transversion-only filtering in this study.

### Selection of windows for divergence-time estimation

We restricted divergence-time analyses to windows with strong phylogenetic support, limited topological discordance, and relatively low among-lineage rate variation. Discordant gene histories can bias branch-length and divergence-time estimates when analysed under a single species-tree topology (Angelis and Dos Reis 2015; Mendes and Hahn 2016), while strong among-lineage rate variation can reduce phylogenetic reliability (Doyle et al. 2015; Vankan et al. 2022).

We applied three filters jointly. First, we required an unrooted Robinson–Foulds distance ≤2 between each window tree and its corresponding species tree (Robinson and Foulds 1981). Second, we required mean ultrafast bootstrap support ≥90 across internal branches. Third, we required the coefficient of variation in root-to-tip branch lengths following midpoint rooting to be ≤0.1 (Vankan et al. 2022).

We concatenated windows that passed all three filters into a single unpartitioned dating alignment and removed columns containing only missing data.

### Relative divergence-time estimation

We estimated relative divergence times using MCMCtree in PAML v4.10.10 (Yang 2007). We used the approximate-likelihood approach of Thorne et al. (1998) and dos Reis and Yang (2011) (Thorne et al. 1998; dos Reis and Yang 2011). We first ran usedata = 3 to estimate branch lengths and calculate the gradient and Hessian, followed by MCMC sampling of divergence times using the resulting likelihood approximation with usedata = 2.

We used the independent-rates relaxed-clock model (clock = 2) (Rannala and Yang 2007), GTR (model = 7), and gamma-distributed among-site rate heterogeneity with six categories. We discarded the first 10^6^ generations as burn-in and sampled 10^7^ subsequent generations.

We aimed to test whether reference choice altered the relative temporal structure of the phylogeny rather than estimate absolute divergence times. We therefore applied no fossil calibrations and fixed the root age to 1, expressing all internal node ages as relative to the root. This removed external calibration as a source of variation among treatments.

We ran two independent MCMC chains for each treatment using different random seeds and the same approximate-likelihood Hessian. Each chain produced 10,000 posterior samples. We pooled the 20,000 samples from both chains to calculate treatment-level posterior mean node ages and 95% highest posterior density intervals, defined as the shortest intervals containing 95% of the pooled samples. We retained the chain-specific posterior means separately to assess variation between independent MCMC runs.

### Comparison among analytical treatments

We compared phylogenetic topology, concordance factors, and relative node ages across the six analytical treatments. For each node, we quantified variation among treatments as the difference between the highest and lowest posterior mean ages calculated from the pooled MCMC samples. We compared this variation with differences between the independent chain-specific posterior means and the widths of the pooled 95% highest posterior density intervals.

To compare missing sequence between read mapping and whole-genome alignment, we restricted all analyses to the same genomic regions. We therefore recalculated missingness for the giant-panda-mapped consensus sequences using the same 24 scaffolds ≥1 Mb used as the whole-genome-alignment anchor.

## Software and data availability

Sequencing accessions are provided in Table S1, and genome-assembly accessions and characteristics are provided in Table S2. PhyloSlide, together with scripts for the LAST and minimap2 whole-genome-alignment workflows used in this study, is available at https://github.com/BiodiversityExtinction/PhyloSlide.

## Acknowledgements

We thank the Novo Nordisk Foundation for supporting M.V.W. through Emerging Investigator grant NNF24SA0093839 and D.A.D. through Emerging Investigator grant NNF23OC0084647. We also acknowledge the use of Claude Opus 5.5 (Anthropic) to assist with code development and manuscript drafting, and ChatGPT GPT-6 (OpenAI) to assist with manuscript drafting and editing. Full responsibility for the analyses, interpretation of results, and final manuscript remains with the authors.

