## Supplementary information for "Reference mapping and whole-genome alignment yield comparable phylogenomic inferences"

**Supplementary tables**

**Table S1.** Sequencing samples and read-mapping statistics used in the reference-genome benchmark. Sample metadata and mapping statistics for the nine ursid datasets included in the study. Each sample was independently mapped to four shared reference genomes: brown bear (*Ursus arctos*), American black bear (*U. americanus*), spectacled bear (*Tremarctos ornatus*), and giant panda (*Ailuropoda melanoleuca*). Where available, each sample was additionally mapped to a conspecific genome assembly.

**Table S2.** Reference genome assemblies used for read mapping and whole-genome alignment.

**Table S3.** Numbers of genomic windows retained at successive stages of the PhyloSlide analyses for each of the four reference-mapping and two whole-genome-alignment (WGA) treatments. No windows were excluded for having fewer than 20 parsimony-informative sites.
